# Analysis of mutations in West Australian populations of *Blumeria graminis* f. sp. *hordei CYP51* conferring resistance to DMI fungicides

**DOI:** 10.1101/696906

**Authors:** M. A. Tucker, F. Lopez-Ruiz, H. J. Cools, J. G. L. Mullins, K. Jayasena, R. P. Oliver

## Abstract

Powdery mildew caused by *Blumeria graminis* f. sp. *hordei* (*Bgh*) is a constant threat to barley production but is generally well controlled through combinations of host genetics and fungicides. An epidemic of barley powdery mildew was observed from 2007 to 2013 in the West Australian wheatbelt (WA). We collected isolates, examined their sensitivity to demethylation inhibitor (DMI) fungicides and sequenced the Cyp51B target gene. Five amino acid substitutions were found of which four were novel. A clear association was established between combinations of mutations and altered levels of resistance to DMIs. The most resistant genotypes increased in prevalence from 0 in 2009 to 16% in 2010 and 90% in 2011. Yeast strains expressing the *Bgh* Cyp51 genotypes replicated the altered sensitivity to various DMIs and these results were confirmed by *in silico* protein docking studies.

## Introduction

*Blumeria graminis* f. sp. *hordei* (*Bgh*) is an ascomycete fungus causing barley (*Hordeum vulgare L*.) powdery mildew. In conducive seasons this biotrophic pathogen can reduce yields by as much as 20% (Murray & Brennan, 2010) but is generally well controlled by host genetics including the durable recessive *mlo* gene (Buschges et al., 1997), dominant major R-genes and combinations of minor genes. In cases where the genetics is inadequate, fungicides can be used. Many classes of fungicides have been used to control powdery mildew but the pathogens have a marked propensity to develop resistance rapidly (FRAG, 2014, Grimmer et al., 2015).

Since 1995 the majority of the West Australian (WA) barley area has been planted to susceptible cultivars and there has been a steep increase in fungicide use (Tucker, 2015). In 2009, 85% of barley crops were treated with one or more fungicide (both seed and foliar) taken from a list of registered chemicals consisting of almost exclusively of DMIs (Murray & Brennan, 2010, ABARES, 2014).

DMI fungicides have been in the forefront of control of fungal pathogens of humans, animals and plants for nearly 50 years (Brent & Hollomon, 2007). These fungicides interrupt the biosynthesis of ergosterol (and other mycosterols in powdery mildews) by inhibiting cytochrome P450 14α-sterol demethylase (*CYP51*) (Senior et al., 1995, Dupont et al., 2012). Resistance is now common in human pathogens, including *Candida* spp. (Xiang et al., 2013, Hull et al., 2012) and *Aspergillus fumigatus* (Lelièvre et al., 2013), and is a serious problem in agricultural systems (Cools et al., 2013, Oliver & Hewitt, 2014). Fungicide resistance has been associated with a number of mechanisms including the alteration and overexpression of the CYP51 as well as enhanced DMI efflux (Omrane et al., 2015, Cools & Fraaije, 2008, Cools et al., 2013).

The most commonly observed mechanism of resistance is non-synonymous changes in the gene sequence of Cyp51 (Cools et al., 2013). A large number of non-synonymous changes have been observed in Cyp51A and B genes of various fungal pathogens. A unified nomenclature for these changes has been proposed and will be adopted in this report (Mair et al., 2016). Two earlier studies examining DMI resistance in *Bgh* in Europe identified *CYP51* changes, *Y137F* and *K148Q* (equivalent to Y136F and K147Q) (Délye et al., 1998, Wyand & Brown, 2005). Isolates with only *Y137F* exhibited both low and high levels of triadimenol resistance and *K148Q* was only ever found in combination with *Y137F*. Hence the exact sensitivity shift afforded by each mutation remains unclear.

West Australian farmers reported a reduction in the effectiveness of DMIs in controlling barley powdery mildew outbreaks since 2005 (GRDC, 2012). Tebuconazole-containing formulations were registered for barley mildew from 1995 in WA (APVMA, 2014) and initially provided good control of leaf rust, powdery mildew and other diseases (Tucker, 2015). However, since 2005, accounts of mildew infection on barley treated with tebuconazole formulations in particular have extended over much of the southern agricultural cropping region with the frequency of reports greatly increasing since 2009 (Lord, 2014).

In this study we have determined the sensitivity of Australian *Bgh* isolates to DMI fungicides registered in WA for use on barley. Sequencing of the *CYP51* coding region in a subset of isolates revealed five mutational changes defining four unique genotypes. The fungicide sensitivities of isolates representing each genotype were determined both by screening on fungicide-treated detached leaves and heterologous expression in *Saccharomyces cerevisiae*. Our results link variations in DMI sensitivity to changes in *CYP51. In silico* protein structural modelling demonstrated the conformational changes afforded by mutations having significant effects on DMI sensitivity and was able to rationalise our observations of partial cross-resistance. A brief report on some this data has been published earlier (Tucker et al., 2015).

## Materials and Methods

### Blumeria graminis f. sp. hordei

#### Isolates

One hundred and nineteen *Bgh* isolates were collected from 2009 to 2013 (Figure 1, Supporting Information Table S1). Isolates from Wagga Wagga, Tamworth (New South Wales) and Launceston (Tasmania) were supplied by the Department of Environment and Primary Industries, Victoria. Isolate purification, sub-culturing and assessments of growth were performed according to Tucker (2013).

**Fig. 1:**
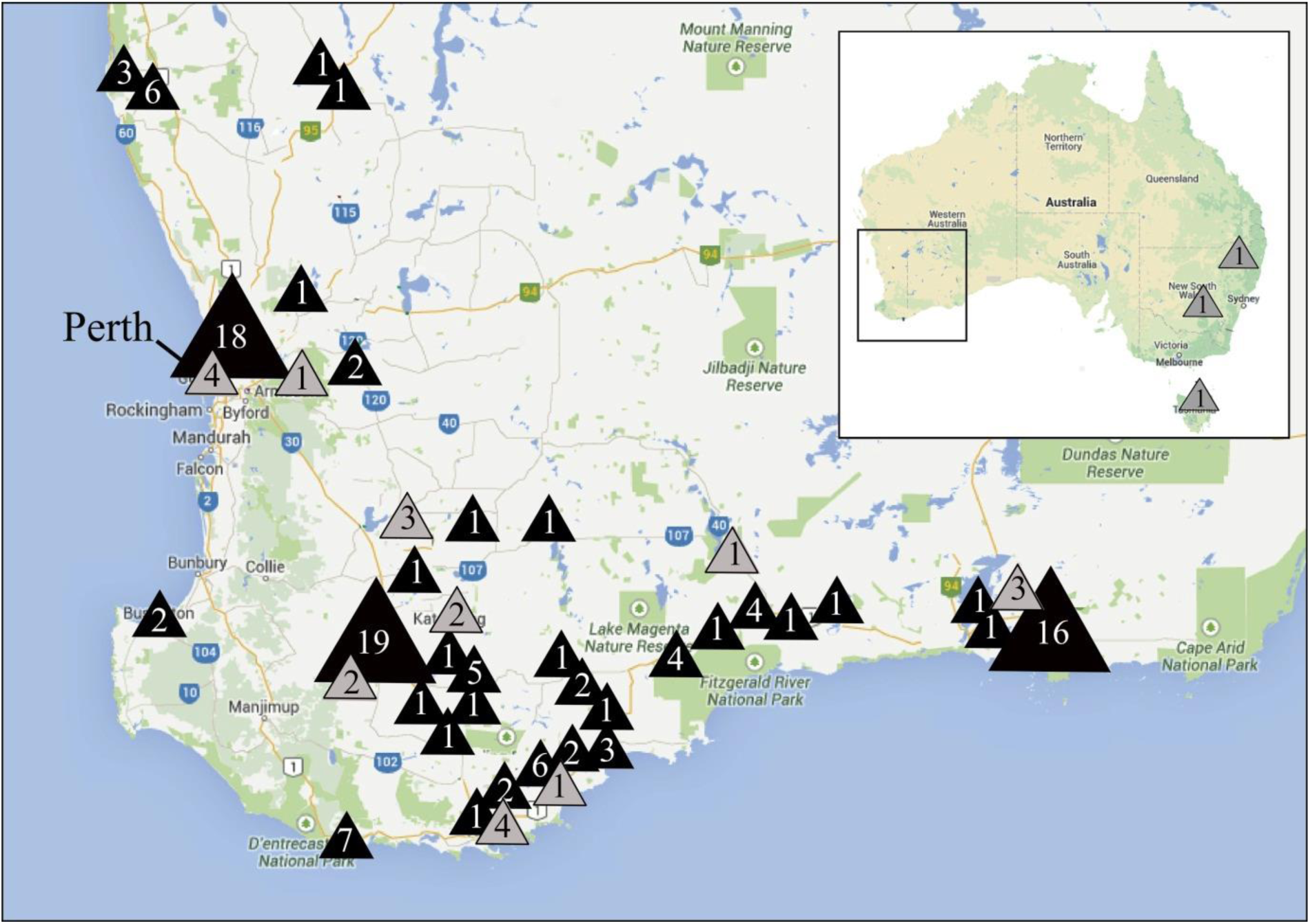
Sample sites of *Bgh* isolates collected from Australia. Black triangles indicate mutant *T524 CYP51* isolates (n = 119). Grey triangles indicate isolates with *CYP51* genotype *S524* (n = 24). Numbers within triangles indicate isolates collected at each site.

**Figure 2.**
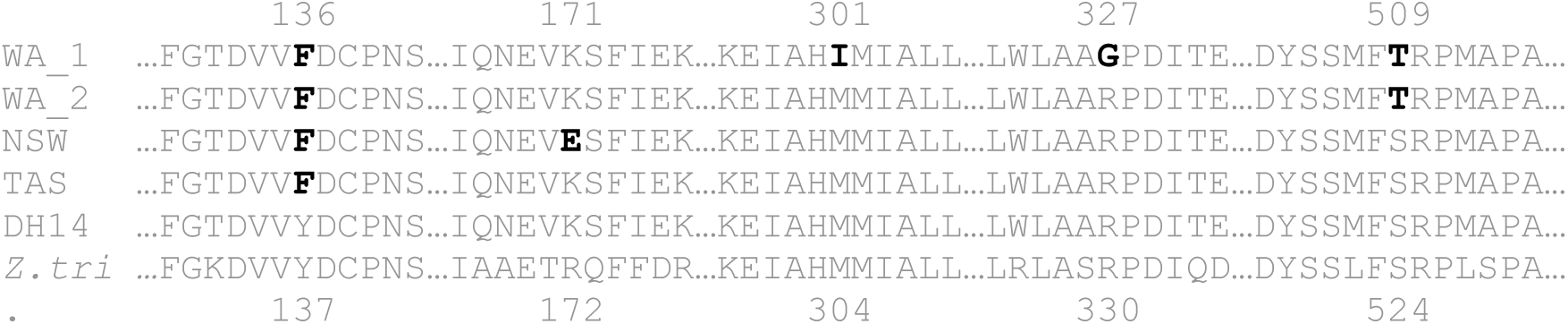
Sequence alignment of fragments of the *Cyp51* protein family. Changes found in Australian *Bgh* isolates to that of the wild-type DH14 (ABSB01000011.1) are indicated in black. Numbers represent amino acid positions. *Blumeria graminis* f. sp. *hordei* isolates WA_1 (Western Australia 1), WA_2 (Western Australia 2), NSW (Wagga Wagga) and TAS (Launceston) are aligned with *Zymoseptoria tritici* isolate ST1 (GenBank Accession AY730587).

#### Fungicide sensitivity assays

Fungicide sensitivities were determined by assessing growth of isolates on susceptible (cv. Baudin) barley leaves inserted into fungicide-amended media. Commercial formulations of DMIs currently registered for *Bgh* control – Laguna (720g L^-1^ tebuconazole, Sipcam), Flutriafol (250g L^-1^ flutriafol, Imtrade Australia), Opus (125g L^-1^ epoxiconazole, Nufarm), Alto (100g L^-1^ cyproconazole, Nufarm), Tilt (418g L^-1^ propiconazole, Syngenta), Proline (410g L^-1^ prothioconazole, Bayer Crop Science), Triad 125 (125g L^-1^ triadimefon, Farmoz) and Jockey Stayer (167g L^-1^ fluquinconazole, Bayer Crop Science) - were incorporated into agar amended with 50mg L^-1^ of benzimidazole (Chan & Boyd, 1992). Middle sections of 10 day old seedlings were excised with each tip inserted abaxial side up into fungicide amended agar. A wide range of concentrations was tested to identify a specific set of 6 needed to calculate an accurate EC50 for each product. Each isolate was inoculated onto three replicates on successive weeks with conidia dislodged 24h before use to promote fresh growth. Conidia were collected on glossy black paper and blown into a 1.5m infection tower to ensure even inoculation. Following 7 days growth at 20±2°C in a 12:12 h light:dark photoperiod, the growth of each isolate was assessed using a 0-4 infection type (IT) scale adapted from Czembor (2000). Each pustule formation was assigned an IT and the average for each isolate and fungicide concentration was determined. Both the average IT and concentration was log transformed, % inhibition calculated and plotted to determine the regression equation and correlation coefficient. The mean 50% effective concentration (EC_50_) with associated errors were calculated for each *Bgh Cyp51* genotype (Fig 3). Data analysis was conducted in JMP, v10 (SAS Institute Inc. Cary, NC).

**Fig. 3:**
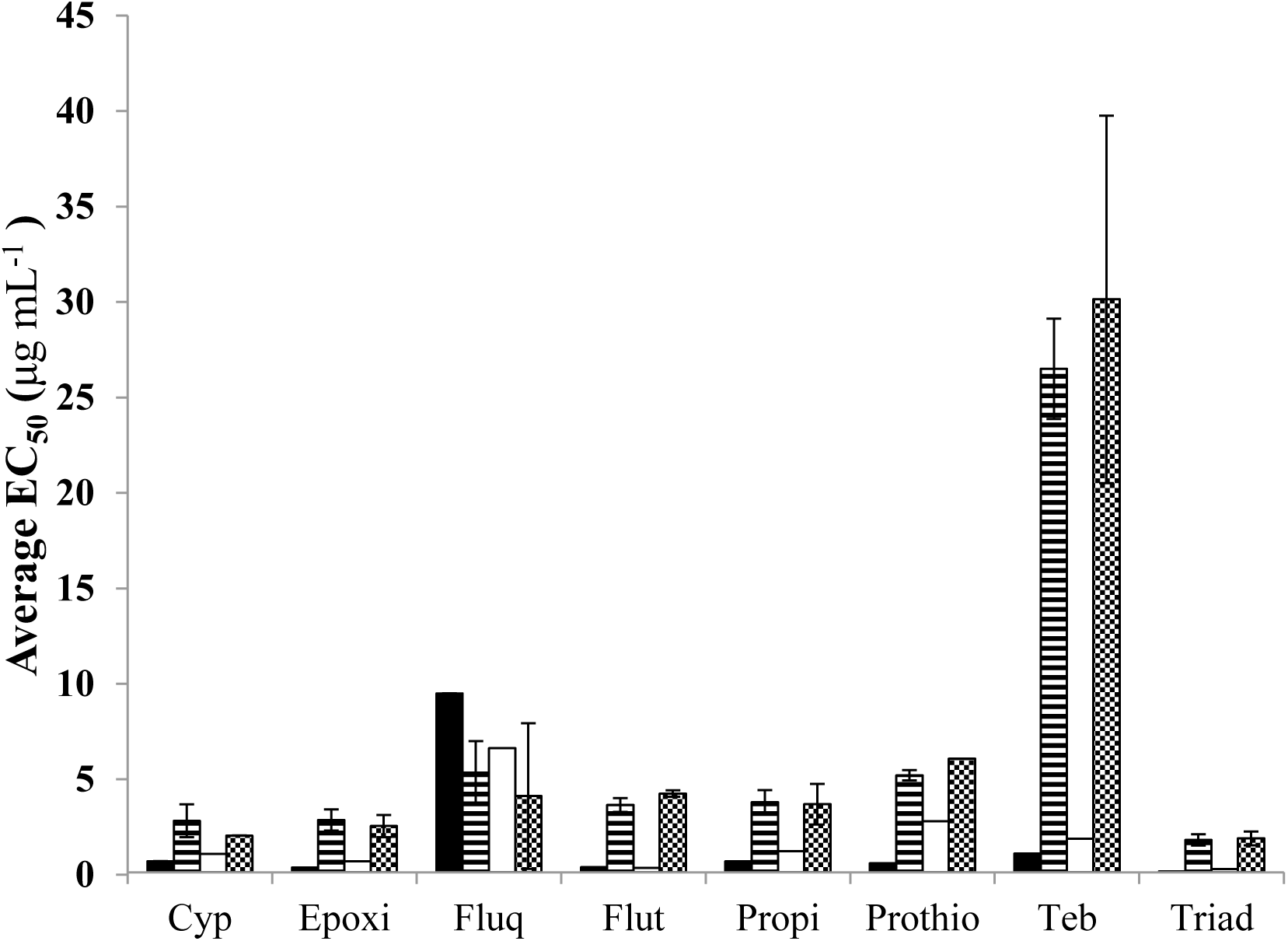
Box plots of the EC_50_ (μg mL^-1^) of a collection of *Bgh* isolates having one of four *Cyp51* genotypes identified in Australia. Genotype 1 (black) - *F137*, genotype 2 – (stripes) *F137/T524*, genotype 3 – (blank) *F137/E172* and genotype 4 – (crosshatch) *F137/I304/R330/T524*. Bars represent mean EC_50_ (μg mL^-1^) of genotypes, with error bars indicated.

#### *CYP51* sequencing

DNA isolations were performed using a BioSprint 15 DNA Plant Kit (Qiagen) following the manufacturer’s instructions. The wild type *Bgh* DH14 isolate (GenBank accession no. AJ313157) was used to design primers (Supporting Information Table S2) covering the entire length of the *Bgh CYP51* (*Bgh51*) gene including the promoter region (Supporting Information Fig. S1). The amplimers of 76 isolates were sequenced using Sanger sequencing and aligned in Geneious v 5.5 (Biomatters). All sequences have been submitted to GenBank (Accession no. KM016902, KM016903, KM016904 and KM016905). A high-throughput method of *S524* and *T524* allele detection was devised ((digesting the amplicon of *Bgh51*_3F and *Bgh51_*3R with Hpy8I) (Supporting Information Table S2)), and used to determine the *CYP51 524* genotype of all 119 isolates.

### Yeast Phenotyping

#### Strains and complementation of transformants

Synthesis of the wild type DH14 (Accession no. AJ313157) *CYP51* gene (*Bgh51wt*) was carried out by GENEWIZ Inc. (South Plainfield, NJ). Terminal restriction enzyme recognition sites for Kpn1 and EcoR1 were added at the 5’ and 3’ ends respectively. The pYES-*Bgh51wt* expression plasmid was constructed by cloning *Bgh51wt* into pYES3/CT (Invitrogen, Carlsbad, CA). pYES-*Bgh51wt* was sequenced to ensure the fidelity and transformed into *S. cerevisiae* YUG37:*erg11* (*MAT*a *ura3-52 trp1-63 LEU2::tTa tetO-CYC1::ERG11*) with its native *Cyp51* gene under the control of a *tetO-CYC1*promoter, repressed in the presence of doxycycline (Parker et al., 2008). All complementation assays were preformed according to Cools *et al* (2010) with photographs taken following 72h of growth at 20°C (Supporting Information Fig. S2). Mutations found in *Bgh51* of Australian isolates were introduced into pYES-*Bgh51wt* through a QuickChange II site-directed mutagenesis kit (Stratagene, La Jolla, CA).

#### Comparative growth rate assay of transformants

The growth rate of transformants was assessed using the Gen 5 data analysis software (BioTek Instruments, Inc. Winooski, VT) where duplicate cultures of replicate transformants were grown in SD GAL + RAF medium (SD medium) overnight at 30°C. One hundred microliters of each overnight culture, at 10^5^ cells ml^-1^, was used to inoculate 3 wells containing 200μl SD medium ±3μg ml^-1^ doxycycline. Cultures were incubated without light at 30°C, and the optical density at 600nm (OD_600_) was measured every 15min for 12 days in a Synergy™ HT Multi-Mode Microplate Reader (BioTek Instruments, Inc Winonski, VT). The mean maximum growth rate for each strain ± doxycycline was determined on the basis of the greatest increase in OD over a 2h period (Supporting Information Table S4).

#### Fungicide sensitivity assays

Sensitivity assays were carried out as described by Cools *et al.* (2010) using pure samples of tebuconazole, cyproconazole, propiconazole, epoxiconazole, fluquinconazole, triadimefon, flutriafol and desthioconazole with a fungicide free control. As prothioconazole must be activated in plant tissue (Parker et al., 2013), desthioconazole was used in all yeast assays.

### Structural modelling

Structural modelling of *Bgh51wt* and mutant forms and ligand docking of epoxiconazole and fluquinconazole was undertaken using an automated homology modelling platform as previously described for *Zymoseptoria tritici* CYP51 (Mullins et al., 2011). The volume of the heme cavity of the wild type and variant protein models was determined using Pocket-Finder (Leeds, UK) based on Ligsite (Hendlich et al., 1997).

## Results

### DNA sequencing

A trial set of *Bgh* isolates were assessed for susceptibility on detached leaves to fungicides then used in WA to control powdery mildew. Substantial variation in resistance was observed. Due to the complexities of the phenotyping assay, we decided to sequence the *CYP51* gene first and then determine fungicide sensitivity of isolates from each genotype class.

Primers were designed spanning both the coding and promoter region of the single *CYP51B* gene (Becher & Wirsel, 2012) in *Bgh* (Supporting Information Table S3, Supporting Information Fig. S1). The *Bgh51wt* DH14 sequence was used as a reference (Spanu et al., 2010). *CYP51* was sequenced from 76 Australian isolates collected from 2009 to 2013, including three from Eastern Australia. No indels were found in the promoter but two synonymous and five non-synonymous changes were identified in the coding region (Fig. 2). All Australian isolates carried previously seen the tyrosine to phenylalanine mutation at amino acid position 136 (*Y137F*) (Wyand & Brown, 2005, Délye et al., 1998). All three isolates from the east of Australia carried two synonymous changes at nucleotides 81 and 1475, which were absent in WA isolates. Further non-synonymous mutations were found; *K172E* (K171E), *M304I* (M301I), *R330G* (R327G) and *S524T* (S509T) in various combinations (Figure 2). Considering only the non-synonymous changes, four novel *Bgh51* genotypes were distinguished. Isolates collected in WA were either *F137/T524* (genotype 2) or *F137/I304/G330/T524* (genotype 4) while isolates from other Australian states were either *F137/E172* (genotype 3) or *F137* (genotype 1). Mutations *I304* and *G330* were consistently found together in the same isolates (Figure 2).

There was both spatial and temporal variation in the frequency of genotypes (Figure 1, Supporting Information Table S1). All isolates collected in 2009 were wild type at *CYP51* position 524. The proportion of mutant *T524* isolates dramatically increased over subsequent seasons; 99 of the 116 WA isolates collected in 2011 contained the *T524* mutation. These mutants were found in all major WA barley growing areas (Figure 1).

### DMI sensitivities of *Bgh* isolates

The sensitivities of 18 *Bgh* isolates (2 isolates from genotypes 2 and 3; 7 from 1 and 4) were determined using detached barley leaves inserted into DMI-amended agar. The results varied between genotype and fungicide (Figure 3, Supporting Information Fig. S4). There was no significant differences in the mean EC50 of *S524* isolates (genotypes 1 and 3) or between isolates with the *T524* mutation (genotypes 2 and 4). Isolates of genotypes 2 and 4 were found to have significantly higher mean EC50 values than genotypes 1 and 3 for most of the DMIs tested. The mean EC50s for *T524* genotypes ranged from 1.88 ug.ml^-1^ for triadimefon, 3.73 ug.ml^-1^ for propiconazole to 29.88 ug.ml^-1^ for tebuconazole. The estimated resistance factors ranged from 3.41 for propiconazole to 17.6 for tebuconazole. However, for fluquinconazole (used solely in WA in seed dressing formulations) mutant *T524* genotypes were marginally more sensitive (EC50 4.73 ug.ml^-1^; RF = 0.58). Unfortunately, due to quarantine restrictions we were not able to phenotype the *Bgh CYP51* DH14 isolate carrying the wild type *Y137* allele.

### Heterologous expression in yeast

The *Bgh51wt* gene was synthesized and cloned into *S. cerevisiae* YUG37:*erg*11 with a doxycycline repressible promoter. The *S. cerevisiae Bgh51wt* transformant was able to grow in the presence of doxycycline (Supporting Information Fig. S2) and for most variants there was no significant difference in the growth rates in the absence of doxycycline. Two *S. cerevisiae Bgh51* variants (pYES-*Bgh51*_*Y137F/S524T/R330G* and pYES-*Bgh51*_*Y137F/M304I/R330G/S524T*) had significantly lower rates and were therefore removed from all further *in vitro* analysis.

The DMI sensitivities of *S. cerevisiae* strains expressing *Bgh51* variants which restored growth on doxycycline-amended medium were determined (Supporting Information Table 4) and resistance factors were calculated (Table 1). Modest RFs were associated with the solo *K172E* and *M304I* mutations. RFs for the *S524T* mutation varied from 0.5 for fluquinconazole to 12.4 for propiconazole. The combination of *F137* and *T524* had much larger RFs of 340.6 for propiconazole and 33.2 for tebuconazole. RFs for fluquinconazole were <1.0 except for the solo *Y137F* construct with a calculated RF of 9.7.

**Table 1:** Resistance factors of *S. cerevisiae* YUG37:*erg11* transformants.

| Construct containing<br>mutation/s | Resistance Factors |  |  |  |  |  |  |  |
| --- | --- | --- | --- | --- | --- | --- | --- | --- |
|  | Teb | Epoxi | Propi | Desthio | Cyp | Flut | Triad | Fluquin |
| pYES-*Bgh51_Y137F* | 1.1 | 3.7 | 1.6 | 3.3 | 1.0 | 1.4 | 1.5 | 9.7 |
| pYES-*Bgh51_K172E* | 0.9 | 1.4 | 0.6 | 2.1 | 0.9 | 1.0 | 0.9 | 0.2 |
| pYES-*Bgh51_M304I* | 0.9 | 1.8 | 2.1 | 0.2 | 0.7 | 0.5 | 0.5 | 0.1 |
| pYES-*Bgh51_S524T* | 3.7 | 7.3 | 12.4 | 1.2 | 2.1 | 2.7 | 3.6 | 0.5 |
| pYES-*Bgh51_Y137F/K172E* | 1.3 | 2.4 | 1.9 | 0.9 | 0.9 | 1.3 | 3.3 | 0.2 |
| pYES-*Bgh51_Y137F/S524T* | 33.2 | 18.5 | 340.6 | 0.8 | 3.2 | 10.5 | 23.8 | 0.2 |
| pYES-<br>*Bgh51_Y137F/S524T/M304I* | 2.3 | 4.9 | 2.8 | 0.1 | 0.8 | 1.9 | 5.7 | <0.1 |

Resistance factors (RF) were calculated from the mean EC<sub>50</sub> values of eight independent replicates. RF <1 indicates greater sensitivity than the wild-type construct. No growth was observed for the pYES3/CT construct.

### Structural modelling

Protein variants of all *Bgh51* genotypes were modelled *in silico* (Supporting Information Fig S5). The effect that each mutational change had on the volume of the heme cavity containing the DMI binding site and the morphological changes to the cavity access channel were determined (Table 2). Modest volume increases in binding cavity were observed with the introduction of the solo mutations; a 17.7% increase with *K172E* and 39.6% increase in volume with *Y137F*. Mutation *S524T* was an exception, with an increase in the volume of the heme cavity by 73.2% compared to that of the wild type model. The combination of *F137/T524* gave an even more substantial increase in volume of 83.9%. Table 2 also shows the estimated distances between amino acids *Y226* (Y222) and *S521* (S506), which span the entrance to the channel leading to the DMI binding site. Modelling simulations predicted that all *Bgh* CYP51 mutations observed in WA would cause a restriction in the diameter of the access channel when compared to wild type *Bgh* CYP51. The most dramatic decrease was observed with the introduction of the *Y137F* mutation, which caused a 28.5% decrease in channel diameter compared to the wild type model. The combination of *Y137F* and *S524T* in a single model did not result in a further significant restriction (Table 2).

**Table 2:** Measurements of heme cavity volume and key inter-residue distances in wild-type genotype *Y137*/*K172*/*M304*/*R330*/*S524* (WT) and mutant *Bgh* CYP51.

| CYP51 genotype | Heme<br>cavity<br>volume<br>(Å<sup>3</sup>) | ΔHCV<sup>a</sup><br>from<br>WT | Diameter<br>channel to<br>binding<br>site<sup>b</sup> | ΔChannel<br>diameter<br>from WT | ΔHCV x<br>ΔChannel<br>diameter |
| --- | --- | --- | --- | --- | --- |
| Wild-type | 1809 | - | 12.862 |  |  |
| *Y137F* | 2526 | +39.6% | 9.202 | -28.5% | 0.113 |
| *K172E* | 2130 | +17.7% | 11.861 | -7.8% | 0.014 |
| *M304I* | 2573 | +42.2% | 12.015 | -6.6% | 0.028 |
| *R330G* | 2607 | +44.1% | 12.074 | -6.1% | 0.027 |
| *S524T* | 3134 | +73.2% | 10.233 | -20.4% | 0.149 |
| *Y137F/K172E* | 2334 | +29.0% | 10.870 | -15.5% | 0.045 |
| *Y137F/S524T* | 3327 | +83.9% | 9.294 | -28.7% | 0.241 |
| *Y137F/M304I/R330G/S524T* | 2181 | +20.6% | 9.960 | -22.6% | 0.047 |

<sup>a</sup>ΔHCV – change in heme cavity volume.
<sup>b</sup>Distance between key amino acids Y226-S315 which border the entrance to the DMI binding site.

Further morphological changes were observed that may impact DMI binding. In particular conformation of a loop of beta turn running from *S520* (S505) to *F523* (F508) is markedly different in the *Y137F* genotype to that of the wild type, with the result that it projects into the cavity. A similar constriction is observed for the *F137/T524* mutant (Supporting Information Fig S5b). However, in this case it is also accompanied by a substantial increase in cavity volume (Table 2), consistent with the exceptional resistance factors observed. It is interesting to note that this loop is adjacent to *S524*. This supports the idea that the structural changes brought about by the *Y137F* mutation on its own may exert selective pressure on the 524 position, leading to the *F137/T524* mutant.

Fluquinconazole docking studies were carried out to elucidate the mechanistic reasons for the contrasting cross resistance patterns (Figure 4b). In the wild type structure, the binding site of fluquinconazole is bordered by amino acids *Y123* and *Y226*. It appears that the position of *Y123* is particularly important in establishing the correct orientation of fluquinconazole so as to be coordinated by the heme. This arrangement is disrupted in the *Y137F* mutant, where *Y123* and *S521* prevent fluquinconazole accommodation (Figure 4b). With the *Y137F/S524T* mutant, *Y123* is positioned similarly to the wild type, allowing accommodation of fluquinconazole as in the wild type. Here *S521* borders the binding site and is predicted to interact with the fluquinconazole ligand (Figure 4c). Thus it appears the relative inconsistency of the *Y137F* mutant and enhanced selection of the *Y137F/S524T* double mutant can be explained by the 3D docking results.

**Fig. 4:**
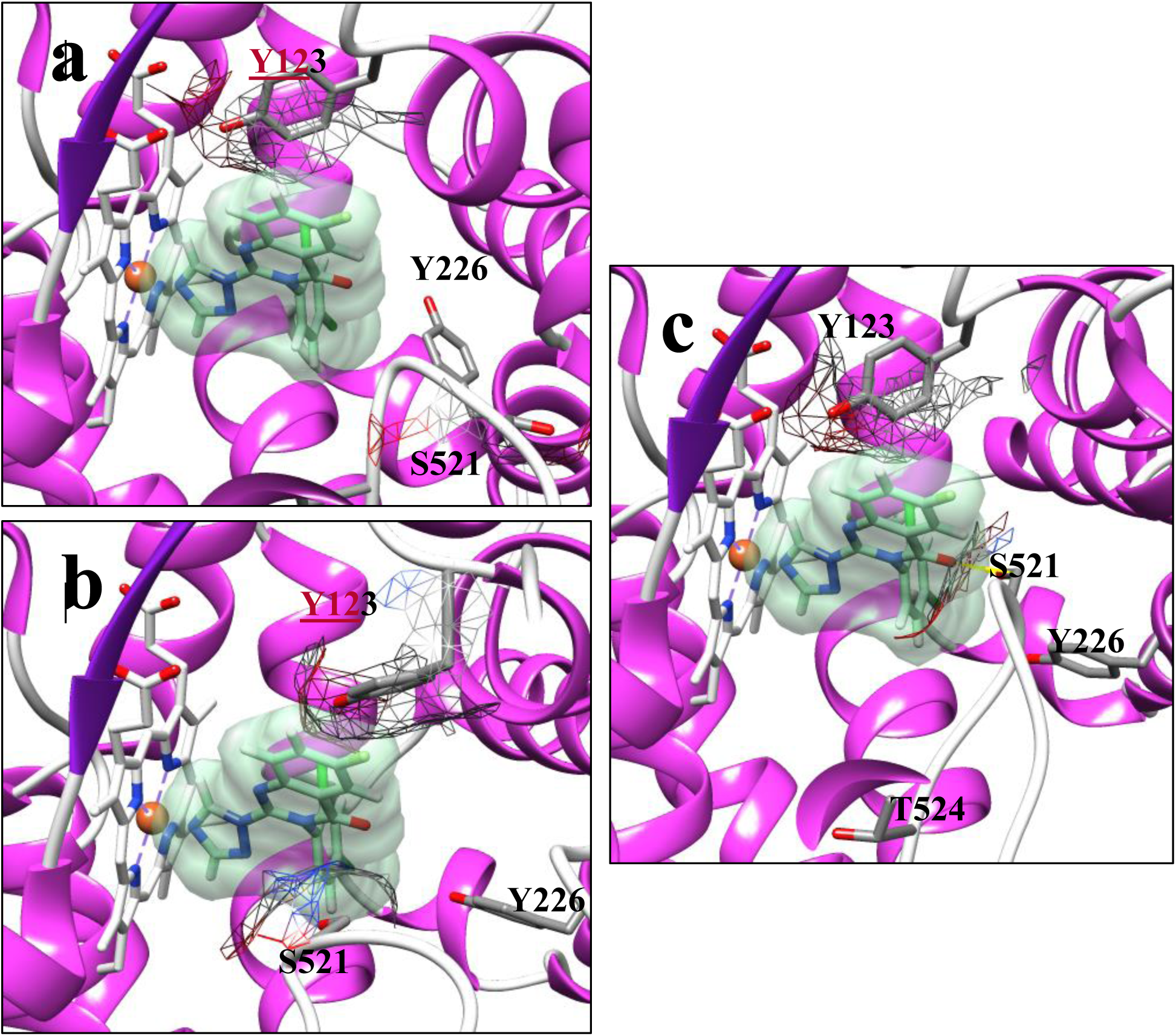
Docking of fluquinconazole in *Blumeria graminis* f. sp. *CYP51*. A) Wild type CYP51, showing bound fluquinconazole (in light green, centre) and steric interaction with *Y123* (surface shown as mesh). B) The *Y137F* mutant, showing encroachment of *Y123* and *S521* (surface shown as mesh) upon the docking site of fluquinconazole, indicating that the compound cannot be bound at that location. C) The *Y137F-S524T* mutant, showing orientation of *Y123* similar to wild type and predicted interaction with S521 (shown in yellow).

## Discussion

Studies best exemplified by the wheat pathogen *Zymoseptoria tritici* have discussed the relationship between mutational changes in *CYP51* with failures of DMI fungicides in the field. DMIs have been used since the first registration in the UK of triadimefon in 1973 UK (Russell, 2005). Twenty years later, *Z. tritici* isolates were found with *CYP51* changes conferring reductions in sensitivity (Cools et al., 2013). Subsequently, numerous DMIs and related compounds have been introduced and 34 additional *CYP51* mutations have been identified. This long history of chemical use and the comparatively recent identification of mutations has made it difficult to discern cause and effect. The situation in WA is far simpler: DMI use has been widespread only since 2004 with the first reports of resistance dating from 2005. Furthermore, usage in WA has been dominated by tebuconazole and propiconazole (Tucker et al., 2015).

Analysis of the single *CYP51* gene of Australian *Bgh* isolates collected from 2009 to 2013 revealed four genotypes. The sensitivities of isolates from different genotypes on detached leaves varied between the DMIs tested. *Bgh* isolates with genotypes harbouring the *S524T* mutation were less sensitive to all the foliar fungicides used on barley in WA and more sensitive to fluquinconazole. The *Y137F* mutation was found in all isolates examined including those from the east of Australia, where as yet there have been no reports of DMI field failure. Previous phenotypic tests (Wyand & Brown, 2005, Délye et al., 1998) correlated the presence of the *Y137F* mutation with strong resistance to triadimenol. We were unable import the wild type *CYP51 Bgh* isolate DH14 in Australia due to quarantine restrictions. However *Y137F* expression in the heterologous yeast system showed only modest decreases in sensitivity to most DMIs including triadimenol (Table 1). This suggests that *Y137F* would lead to only small reductions in the DMI field efficacy. The ubiquity of *Y137F* in Australia suggests two possibilities; (1) the limited fungicide use in eastern of Australia has been sufficient to select for this mutation or (2) the wild type CYP51 genotype has never been present in Australia.

A search was conducted on the *CYP51* mutations in other fungal species reported as conferring a reduction in DMI sensitivity. The *Bgh51* amino acid sequence of Australian genotypes was aligned with *Z. tritici* CYP51 (Figure 2). Mutational changes at the amino acids *137, 304, 330* and *524* fall in regions conserved between *Bgh* and *Z. tritici* (Becher & Wirsel, 2012). Amino acids 136 and 509 in *Bgh51* correspond to 137 and 524 in *Z. tritici* which have previously been correlated with alterations in DMI sensitivity (Cools et al., 2011). The combination of *Y137F* and *S524T* was associated with substantial RFs in both the *Z. tritici* strains and the yeast transformants. This study did not test fluquinconazole or the solo *Y137F* genotype in the yeast system.

In the current study the combination of *Y137F* and *S524T* encoded a CYP51 with a marked decrease in sensitivity to tebuconazole and propiconazole in both the mildew strains and the yeast system. This may be sufficient to account for the field failure (Figure 3). Increases in heme cavity volume and restriction of the access channel in *Y137F/S524T* protein models correlate well with the significant RF obtained (Figure 5). A high RF for propiconazole was also observed for the *Y137F/S524T Bgh CYP51* construct when expressed in the yeast system.

**Fig. 5:**
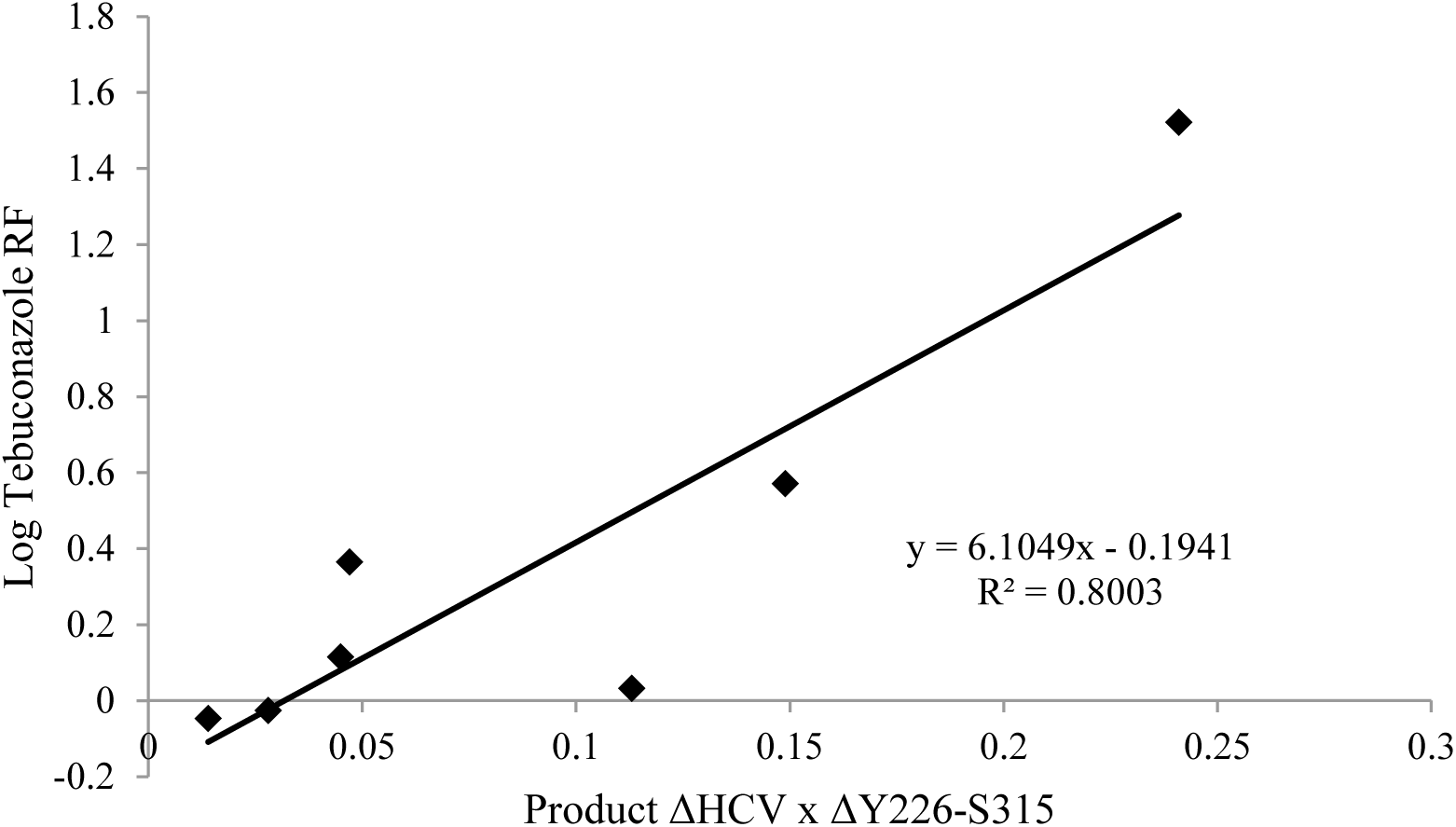
Correlation between tebuconazole resistance factor (RF) of pYES-*Bgh51*_*Y137F*/*S524T* and the product of the change in the volume of the heme cavity (ΔHCV) with the change in distance between amino acids *Y226* and *S315* (ΔY226-S312).

Structural modelling suggests that there are two main mechanisms that underpin the emergence of DMI resistance associated with mutational changes in *Bgh51*. The first mechanism is similar to that observed in *Z. tritici* CYP51 (Mullins et al., 2011), where the gross volume of the heme cavity increases with successive mutations (Table 2). There appears to be a correlation between the increase in cavity volume and the RFs reported in table 1. It is likely that any increase in heme cavity volume would perturb the orientation of the DMI ligand and hence its binding to the heme. This therefore differentiates the smaller DMI ligands such as tebuconazole and epoxiconazole.

The second mechanism at play provides a means of linking structural changes with phenotypic changes in a measurable way. Changes in distances between specific pairs of residues that border the cavity result in changes to the diameter of the access channel. The limiting of the binding surface between *Y226* and *S314* (S312) appears to correlate well with resistance to tebuconazole. The narrowing of the access channel between *Y226* and *S521* correlates particularly well, especially when tempered by consideration of the effects of each variant on the cavity volume. This is demonstrated by the result obtained when the product of the percent change in the heme cavity volume is multiplied by the percent change in the distance between *Y226* and *S521* (Figure 5). All the variants that contain *F137* demonstrate a substantially reduced distance between *Y226* and *S521* (Table 2). When one of the mechanisms is employed, moderate resistance factors are observed (*F137* (access channel narrowing); *T524* (substantial increase in cavity volume)). Although, when both mechanisms act together there is a strong correlation between the structural changes and the very high resistance factors of the *F137/T524* mutants in the presence of tebuconazole. The *in silico* creation of *Bgh51wt* and mutant CYP51 protein variants opens the possibility of future docking studies employing novel or unregistered DMI fungicides. This will allow the prediction the effectiveness of any new product prior to *in planta* testing. Furthermore, we can now recommend bespoke spray regimes depending on which *Bgh51* genotype is present in the field.

One of the major resistance strategies used for fungicides is to mix active compounds with different MOA because isolates with mutations conferring resistance to one fungicide will most likely still be sensitive to the second (Van Den Bosch et al., 2014). This strategy requires that there is no positive cross resistance between the two fungicides and so generally rules out mixtures of the same MOA. However some cases of negative cross-resistance within a single MOA group has been shown with *Z. tritici* isolates which are highly resistant to tebuconazole but fully susceptible to prochloraz (Leroux et al., 2007, Fraaije et al., 2007). The negative cross-resistance shown in both the *Bgh* (Figure 3) and yeast expression studies (Table 1) was confirmed using *in silico* protein docking studies. Here the single *Y137F* mutation substantially impaired the binding of fluquinconazole (Figure 4). In contrast, the binding of fluquinconazole at the docking site of the *Y137F/S524T* protein model was much akin to that of the wild-type.

The widespread use of highly susceptible varieties and the repeated use of a single MOA fungicide was a perfect recipe for an epidemic of fungicide resistance in West Australia. A review covering the decade from 1999 to 2009 estimated that *Bgh* in WA caused losses of AU$30M p.a. (Murray & Brennan, 2010) However, given the spread of the highly virulent (Tucker et al., 2013) and DMI-resistant population of *Bgh* throughout the barley growing regions of WA, the losses can now be estimated to have been about AU$100M p.a from 2007 to 2010 (Tucker, 2015).

## Acknowledgements

We would like to thank Ryan Fowler from the Department of Environment and Primary Industries, Victoria, for supplying isolates. Further thanks to Harry Zhang for his technical assistance and to Simon Ellwood for the supply of isolates.

## Abbreviations

Cyp: Cyproconazole
Desthio: Desthioconazole
Epoxi: Epoxiconazole
Fluquin: Fluquinconazole
Flut: Flutriafol
Propi: Propiconazole
Prothio: Prothioconazole
Teb: Tebuconazole
Triad: Triadimefon

## Supporting Information

**Supp Table S1.** Origin of isolates used in the study.

| Collection year | Collection site | State<sup>a</sup> | Number of isolates<sup>b</sup> | *CYP51* genotype<sup>c</sup> |
| --- | --- | --- | --- | --- |
| 2009 | Beverly | WA | 1 | 1 |
| 2009 | Katanning | WA | 1 | 1 |
| 2009 | Mt Madden | WA | 2 | 1 |
| 2009 | South Perth | WA | 1 | 1 |
| 2009 | Williams | WA | 1 | 1 |
| 2010 | Kojonup | WA | 2 | 1 |
| 2010 | Scaddan | WA | 1 | 2 |
| 2010 | South Perth | WA | 1 | 1 |
| 2010 | South Stirlings | WA | 7 | 1 |
| 2010 | South Stirlings | WA | 1 | 2 |
| 2011 | Albany | WA | 3 | 4 |
| 2011 | Boxwood Hill | WA | 3 | 4 |
| 2011 | Bentley | WA | 2 | 4 |
| 2011 | Esperance | WA | 1 | 1 |
| 2011 | Esperance | WA | 6 | 4 |
| 2011 | Gibson | WA | 5 | 4 |
| 2011 | Gnowellen | WA | 1 | 4 |
| 2011 | Kojaneerup | WA | 3 | 4 |
| 2011 | Northam | WA | 1 | 4 |
| 2011 | South Perth | WA | 1 | 4 |
| 2011 | South Stirlings | WA | 5 | 4 |
| 2011 | Takalarup | WA | 2 | 4 |
| 2011 | Tamworth | NSW | 1 | 3 |
| 2011 | Launceston | TAS | 1 | 1 |
| 2011 | Wagga Wagga | NSW | 1 | 3 |
| 2011 | Wellstead | WA | 2 | 4 |
| 2012 | Badgingarra | WA | 1 | 4 |
| 2012 | Esperance | WA | 1 | 4 |
| 2012 | Esperance | WA | 1 | 1 |
| 2012 | Kojonup | WA | 6 | 4 |
| 2012 | Northam | WA | 1 | 4 |
| 2012 | Southern River | WA | 1 | 4 |
| 2013 | Kojonup | WA | 6 | 4 |

<sup>a</sup>Australian state abbreviations: WA - Western Australia, TAS - Tasmania and NSW - New South Wales.
<sup>b</sup>A high-throughput method was employed to detect the presence of the *S524T* mutation in a further 46 isolates.
<sup>c</sup>Genotypes are as follows 1 –*F137*, 2 – *F137/T524*, 3 –*F137/E172* and 4 – *F137/I304/R330/T524*.

**Supporting Information Table S2:**
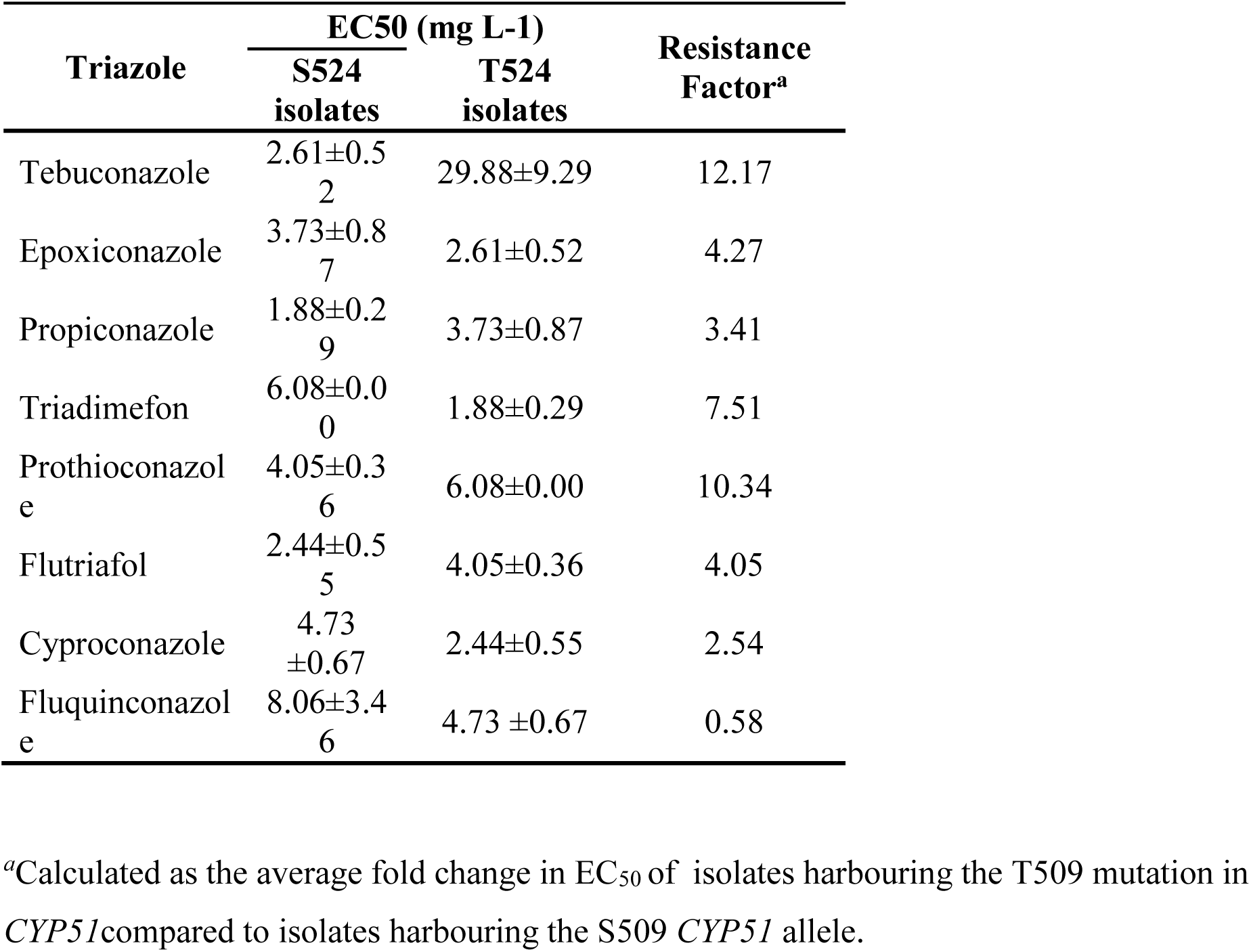
Average *in vitro* EC_50_ and resistance factors of T509 and S509 *Bgh51* isolates when exposed to currently registered triazoles in WA.

**Supporting Information Table S3:**
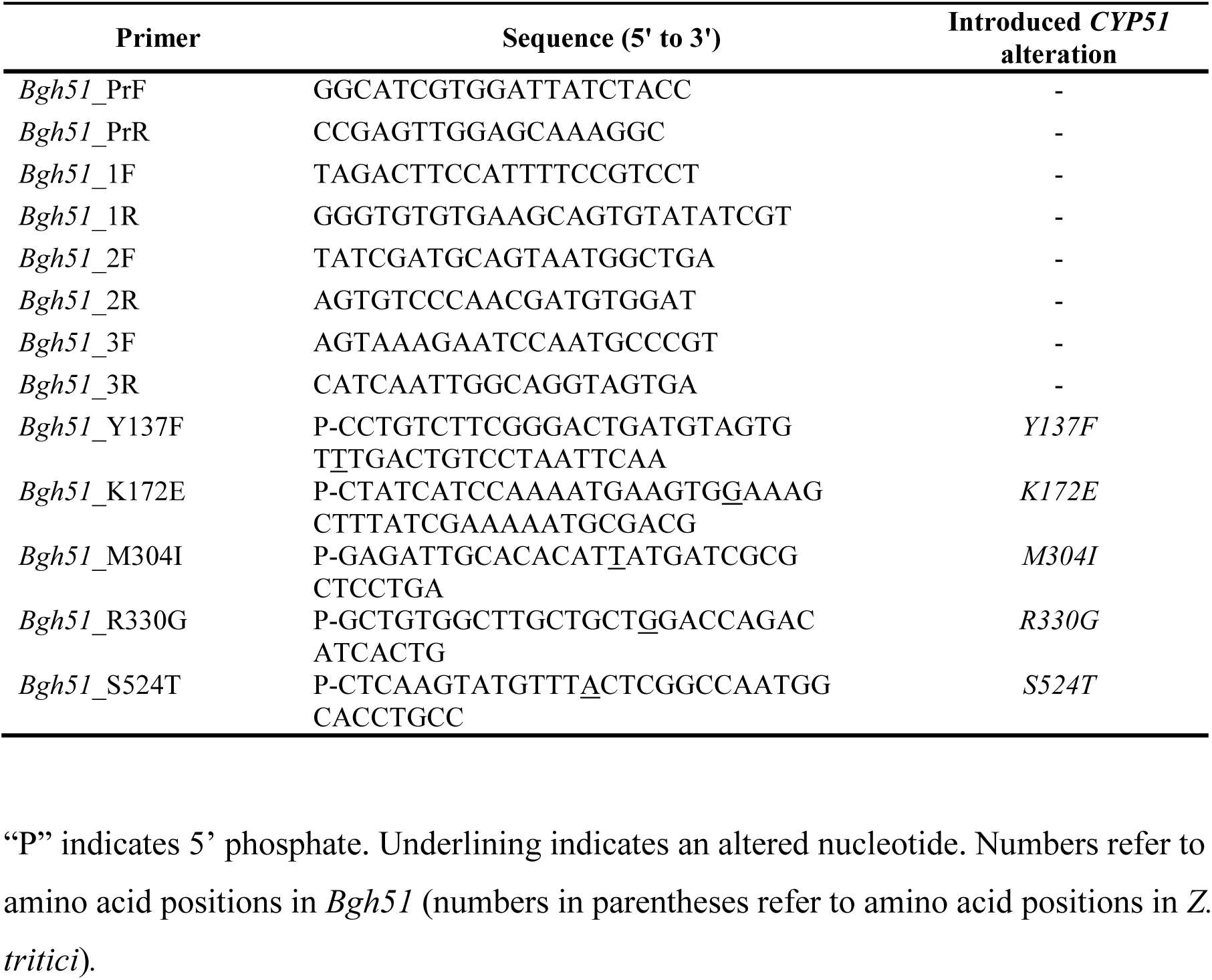
Primers used for sequencing and site-directed mutagenesis.

**Supporting Information Fig. S1:**
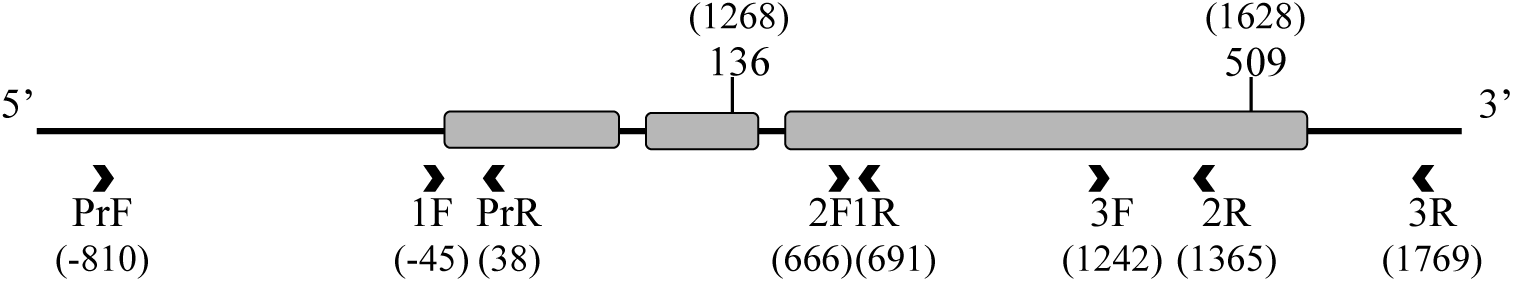
Binding position of primers used to sequence the length of *Bgh51* including the 5’ promoter region. Exons are represented by grey bars with primer binding sites as directional arrows. Names of *Bgh51* primers are in reference to supplementary table 4 with position of 5’ nucleotide given in parenthesis. Positions of amino acids 136 and 509 have been indicated with the position of the mutated base given in parenthesis

**Supporting Information Fig. S2:**
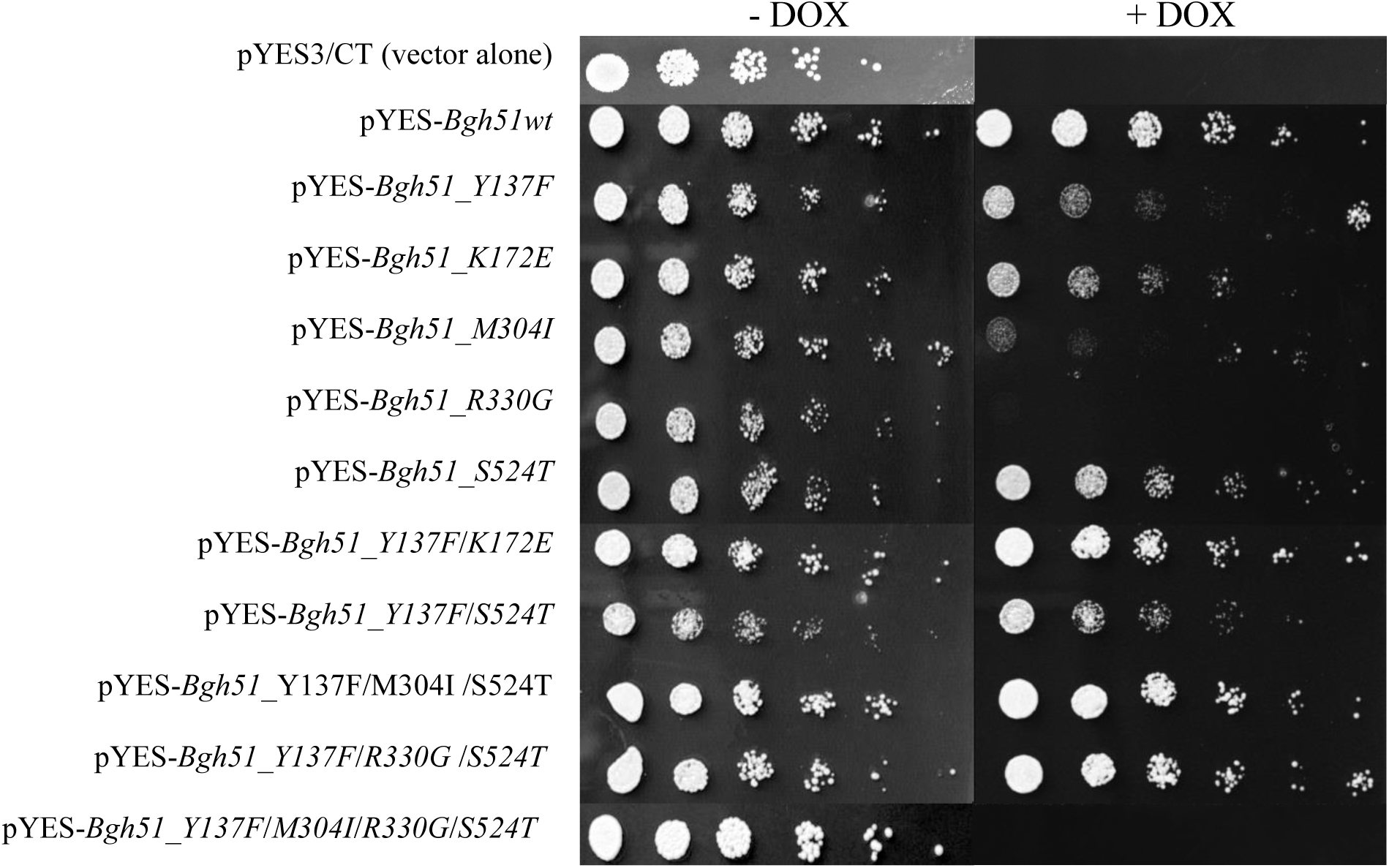
Complementation of *S. cerevisiae* strain YUG37:*erg11* with wild type (*Bgh51wt*) and mutated variants. Growth in the absence (-DOX) and presence (+DOX) of doxycycline is shown. Numbers refer to amino acid positions in *Z. tritici.*

**Supporting Information Table S4:**
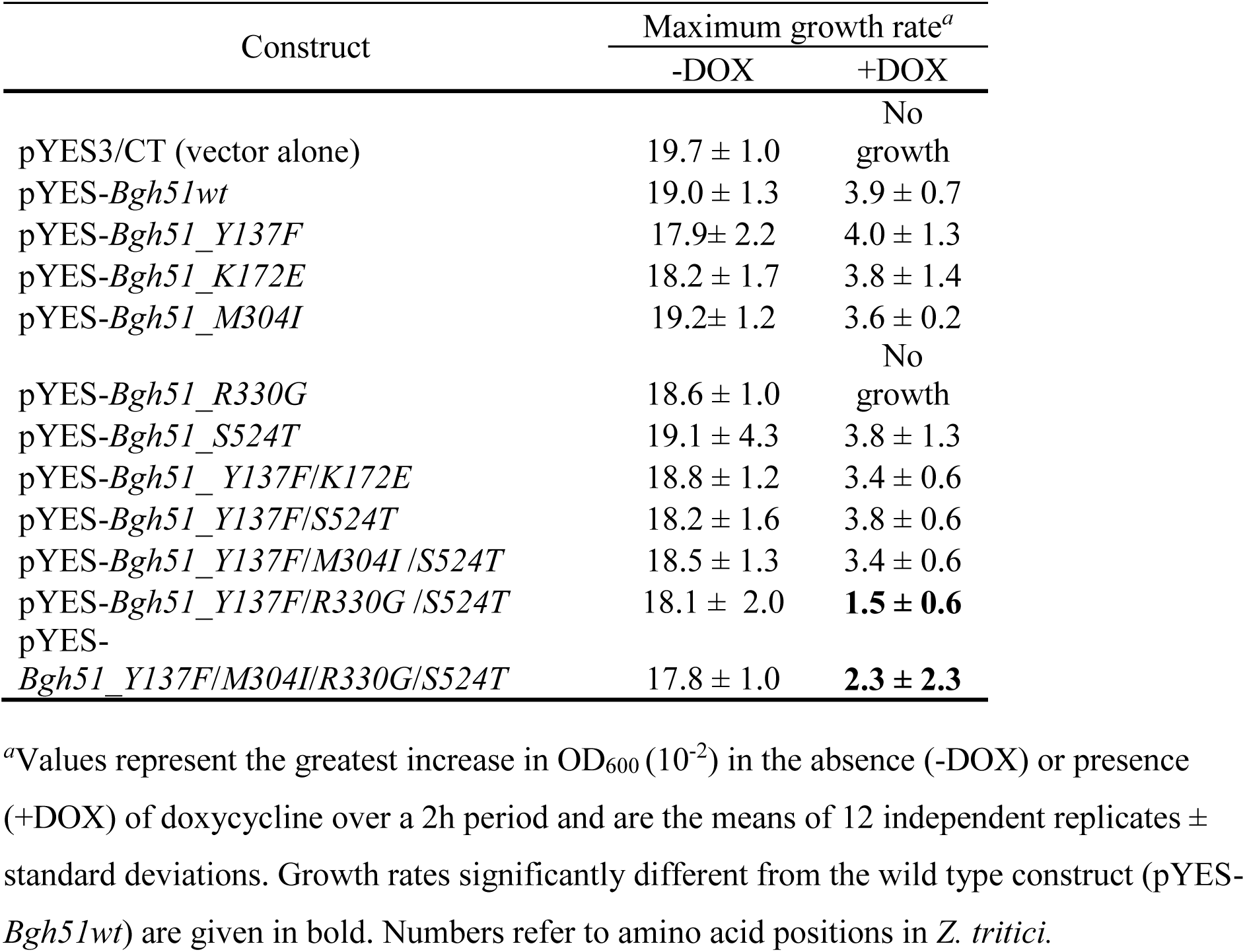
Growth rate analysis of *S. cerevisiae* YUG37:*erg11* transformants.

**Supp Table S5.** *In vitro* EC50 and resistance factors of *T524* (T509) and *S524* (S509) *Bgh* isolates when exposed to currently registered triazoles in WA.

| Fungicide | EC<sub>50</sub> (mg liter<sup>-1</sup>) |  | Resistance Factor |
| --- | --- | --- | --- |
|  | T524 isolates | S524 isolates |  |
| Tebuconazole | 29.88±4.02 | 1.7±0.65 | 17.61 |
| Epoxiconazole | 2.61±0.52 | 0.61±0.48 | 4.27 |
| Propiconazole | 3.73±0.87 | 1.09±0.59 | 3.41 |
| Triadimefon | 1.88±0.29 | 0.59±0.19 | 7.51 |
| Prothioconazole | 6.08±0.00 | 0.59±0.19 | 10.34 |
| Flutriafol | 4.05±0.36 | 0.37±0.26 | 10.92 |
| Cyproconazole | 2.44±0.55 | 0.96±0.64 | 2.54 |

**Supporting Information Fig. S5.**
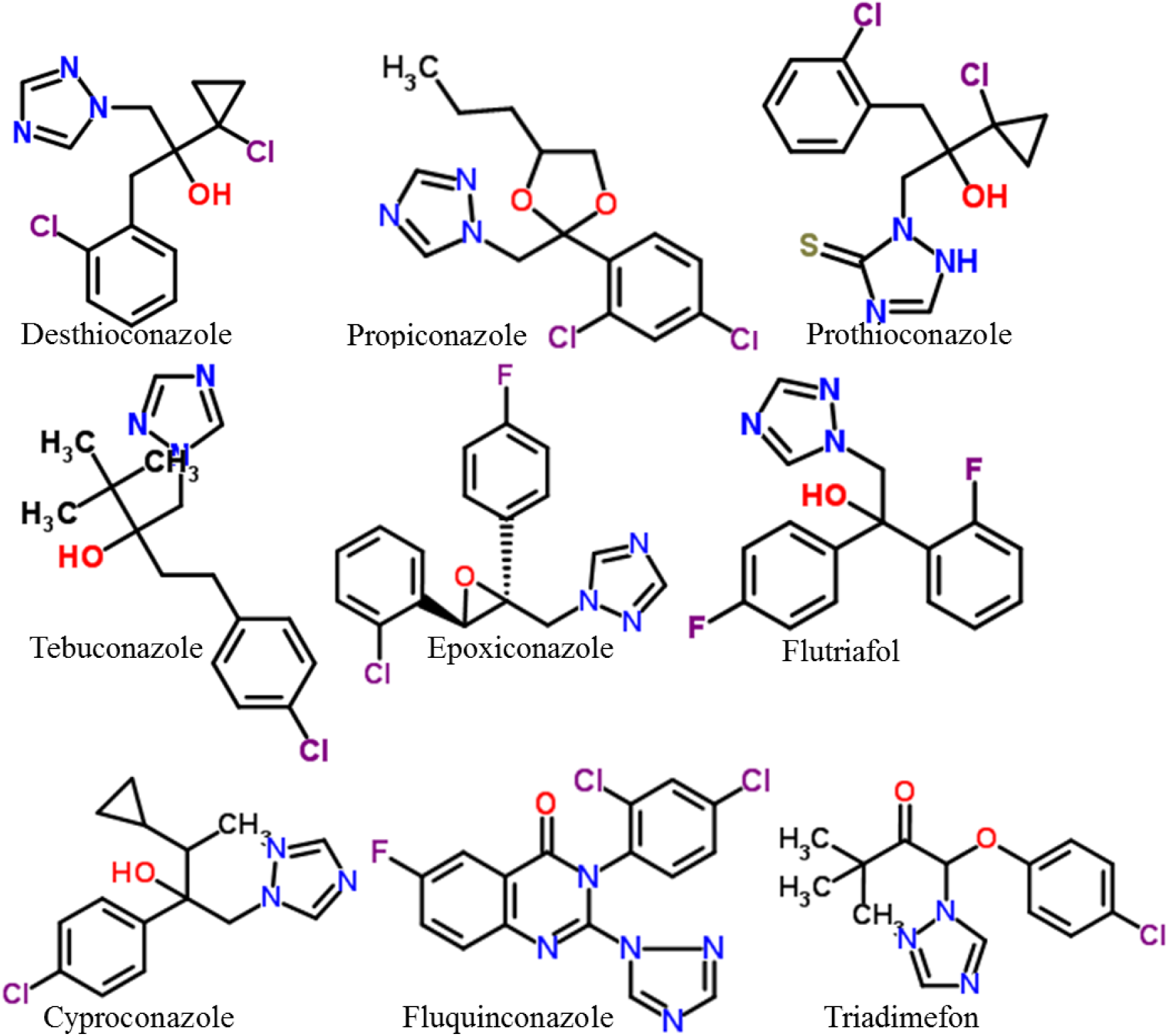
Chemical structure of triazole fungicides employed in this study. Figures used with permission from www.chemspider.com (Accessed 16^th^ September 2015)

